# The interdependent nature of multi-loci associations can be revealed by 4C-Seq

**DOI:** 10.1101/028555

**Authors:** Tingting Jiang, Ramya Raviram, Pedro P. Rocha, Valentina Snetkova, Charlotte Proudhon, Sana Badri, Richard Bonneau, Jane A. Skok, Yuval Kluger

**Author notes:** These authors contributed equally to this manuscript (Jane A. Skok, Yuval Kluger).

## Abstract

Use of low resolution single cell DNA FISH and population based high resolution chromosome conformation capture techniques have highlighted the importance of pairwise chromatin interactions in gene regulation. However, it is unlikely that these associations act in isolation of other interacting partners within the genome. Indeed, the influence of multi-loci interactions in gene control remains something of an enigma as beyond low-resolution DNA FISH we do not have the appropriate tools to analyze these. Here we present a method that uses standard 4C-seq data to identify multi-loci interactions from the same cell. We demonstrate the feasibility of our method using 4C-seq data sets that identify known pairwise interactions involving the *Tcrb* and *Igk* antigen receptor enhancers, in addition to novel tri-loci associations. We further show that enhancer deletions not only interfere with tri-loci interactions in which they participate, but they also disrupt pairwise interactions between other partner enhancers and this disruption is linked to a reduction in their transcriptional output. These findings underscore the functional importance of hubs and provide new insight into chromatin organization as a whole. Our method opens the door for studying multi-loci interactions and their impact on gene regulation in other biological settings.

## INTRODUCTION

In recent years the influence of chromosomal interactions in gene regulation has become increasingly apparent (1-6). Insights into the importance of these associations were initially obtained from studies of at the single cell level using DNA FISH and more recently by molecular analysis using chromosome conformation capture (3C) techniques (7-11). The original 3C assays examined interaction frequencies between two fixed points of interest, while a later iteration of this technique, 4C-seq enabled comparisons of interaction frequencies from a single viewpoint to all other sites across the genome. Hi-C took this a step further providing a tool to analyze all possible pairwise interactions in the nucleus (11). These molecular approaches highlighted the importance of enhancer – promoter interactions in controlling lineage and stage-specific gene expression patterns (12). Moreover these studies demonstrated the significance of chromosome looping in separating out regions that have a distinct chromatin status (13) and the functional relevance of this aspect of chromatin organization in gene regulation (14). Hi-C and 5C further identified the existence of higher-order structures known as topologically associated domains (TADs) that encompass megabase wide stretches of DNA that interact with each other at high frequency (15-18). By definition, DNA contacts occur most frequently within individual TADs, however interactions between TADs on the same chromosome also occur, albeit at a lower frequency, while TADs located on different chromosomes will interact even less often. These analyses have revolutionized our understanding of how structure function connections underpin gene regulation and have provided a platform for future experiments.

One poorly understood aspect of the role of chromatin associations in gene regulation concerns the influence of simultaneous interactions involving more than two partner regions. Although mathematical modeling approaches have attempted to reconstruct three-dimensional structures to address this question, only combined DNA and RNA FISH analyses have unambiguously been able to identify multi-loci interactions and to probe their functional importance (19). However, as mentioned above FISH is low resolution and it is not possible to look at interactions that occur at short length scales between regions of the genome separated by less than a few Kb. Furthermore, FISH approaches are currently largely low-throughput.

Given the potential importance of the synergistic effects of multi-loci interactions on gene regulation it is important to design approaches for detecting and validating these. Ay et al. recently developed an innovative 3C-based technique to detect multi-loci interactions, however, this method has several drawbacks regarding low efficiency of data extraction and low resolution (20). Making use of standard 4C-Seq datasets that assess genome wide spatial contacts from a single viewpoint or bait sequence, we have now established that identification of multi-loci interactions is feasible. We tested our approach using several 4C-seq datasets generated from *ex-vivo* derived sorted developing murine B and T lymphocytes. Here we demonstrate the robustness and reproducibility of our method using four different bait sequences on chromosome 6 associated with the T cell receptor beta (*Tcrb*) locus enhancer, Eβ,and the immunoglobulin kappa (*Igk*) enhancers, MiEκ and 3’Eκ as well as from a region downstream of *Igk* (DSR) located close to the 3’ end of the *Igk* locus. The three antigen receptor enhancers are activated in a lineage and stage specific manner, in line with lineage specific recombination and expression of their associated loci: Eβ is activated in T cells while MiEκ and 3’Eκ are activated in B cells. Using these viewpoints at different stages of lymphocyte development we were able to identify known lineage specific pairwise interactions as well as tri-loci associations involving the bait and two other genomic regions. Specifically, we demonstrate simultaneous association of, MiEκ, 3’Eκ and a third *Igk* enhancer, Edκ in pre-B and immature B cells. Deletion of either the MiEκ or 3’Eκ eradicates this enhancer hub leading to spatial re-organization of neighboring regions that not only interfere with the tri-loci interaction in which the deleted enhancer participates, but also disrupts pairwise associations between the other two enhancers. Moreover, we demonstrate that loss of contact involving these enhancers leads to a disruption of their transcriptional output. Our studies provide a new tool for analyzing gene regulation through identification of multi-loci interactions and for dissecting out hierarchical relationships using 4C-seq.

## MATERIALS AND METHODS

### Cell Sorting

Single cell suspensions were prepared from thymus or bone marrow (BM) from mice of various genotypes. CD4 and CD19 microbeads (Miltenyi Biotec) selections were performed on Thymocytes and BM cell suspension respectively using a Manual MACS^®^ Cell Separator (Miltenyi Biotec) to reduce cell sorting time. Flow cytometry cell sorting was performed on Beckman Coulter MoFlo and Sony SY3200 machines as follow: from CD4^−^ selection, DN2/3 cells were purified as Thy1.2^+^TCRb^lo^CD4^−^CD8^−^ CD25^+^, from CD4^+^ selection DP cells were purified as Thy1.2^+^TCRb^int^CD4^+^CD8^+^ and from CD19^+^ selection, pre-B cells were purified as IgD^−^B220^+^IgM^−^c-kit^−^CD25^+^ and immature-B cells as IgD^−^B220^+^IgM^+^. Antibodies used were as follows: Thy1.2 PE-Cy7 (clone 53-2.1, eBioscience, 1:5,000 dilution), TCR-b APC-eFluor780 (clone H57-597, eBioscience, 1:500 dilution), CD4 APC (clone RM4-5, BD Biosciences, 1:500 dilution), CD8a FITC (clone 53-6.7, BD Biosciences, 1:500 dilution), CD25 PE (clone PC61, BD Biosciences, 1:500 dilution), IgD APC-Cy7 (clone 11-26c.2a, BioLegend, 1:500 dilution), CD45R/B220 PE-Cy7 (clone RA3-6B2, BD Biosciences, 1:500 dilution), IgM FITC (clone II/41, BD Biosciences, 1:500 dilution), CD117/c-kit APC (clone 2B8, eBioscience, 1:500 dilution).

### 4C library generation

4C-seq uses 3C (chromosome conformation capture) products as a template for inverse PCR to amplify the genomic regions interacting with a region of interest, the bait. The 3C template is produced by two successive rounds of digestion-ligation using two different 4bp cutters, NlaIII and DpnII as outlined in **Figure 1A-G**. The first step involves crosslinking the DNA with formaldehyde prior to the first round of digestion-ligation that joins DNA regions in close physical proximity. The second digestion-ligation event occurs following reversal of the crosslinks. This produces small circular fragments for efficient PCR amplification using primers specific for bait regions. Here we generated 4C-seq libraries using bait sequences associated with the *Tcrb* enhancer, Eβ, and *Igk* enhancers, MiEκ and 3’Eκ, as well as a region downstream of *Igk* (DSR). (**Figure 2A**). Illumina-specified adapters for Illumina sequencing were included at the 5’ end of each primer. Our 4C libraries were sequenced on the Illumina Hi-Seq 2000 using single-read 100-cycle runs.

**Figure 1:**
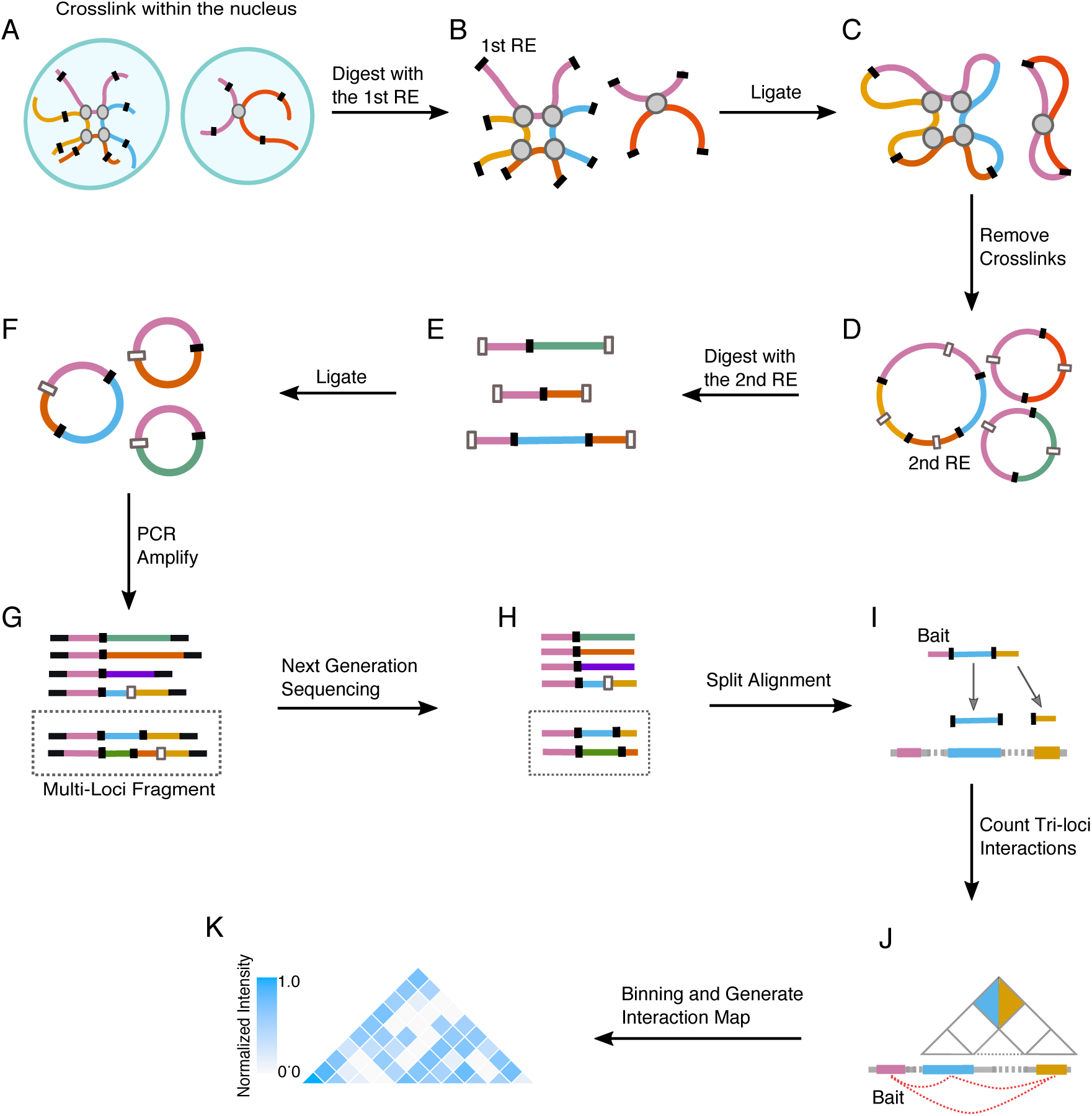
Workflow for extracting multi-loci chromatin interactions using 4C-seq data. **A.** Cross-linking step. **B.** Primary restriction enzyme digestion. **C.** First ligation step. **D.** Removal of crosslinks. **E.** Secondary restriction enzyme digestion. **F.** Second ligation step. **G.** PCR amplification of fragments (tri-loci fragments are shown in the dotted box). **H.** High throughput sequencing (tri-loci reads are shown in the dotted box). **I.** Alignment of tri-loci reads. Each read is split into three segments at primary restriction enzyme cut sites that include the bait segment. These segments are aligned independently to the genome. **J.** Tri-loci interactions along the genome (bottom) and their representation in a heatmap (colored squares indicate simultaneous interaction of the blue and orange genomic locations with the bait). **K.** Heatmaps were generated by binning the genomic region of interest in a resolution dictated by the genomic length of the loci involved in the study.

At least two replicates were processed for each sample. The number of reads obtained for each dataset is listed in **Supplementary Table 1**.

### Alignment of reads representing multi-loci interactions

For each single-end read, the bait sequence was removed from the 5’ end. The remaining part of each read was split as shown in **Figure 1H**. Prior to alignment we extended the first interacting segment with CATG sequences (the NlaIII restriction enzyme cut site) upstream and downstream of this string. Further, we extended the second interacting segment with one CATG sequence upstream of the string. Each string of reads with multiple fragments, was mapped individually to the mouse reference genome (mm10) using Bowtie (1.01) with parameters as follows:

-S -v 2 -k 1 -m 1 -I 0 --best --strata. Alignment of segments was sorted by read name using samtools (0.1.18). We further removed pairs of interacting segments that align to a contiguous region in the genome. These pairs represent only one interacting locus that contains an undigested CATG site. The remaining reads with two segments aligned to two different genomic locations were termed ‘tri-loci’ fragments (**Figure 1I**). These interactions are represented as a heatmap (**Figure 1J**)

### Reducing the effect of PCR artifacts in Multi-loci interactions

To take account of PCR artifacts that can be generated during the amplification step, we created a histogram whose y-axis represents the number of different tri-loci interactions as a function of the number of identical PCR duplicates or real interacting loci shown in the x-axis (**Figure S1**). To reduce the effect of PCR artifacts we capped the number of tri-loci fragments above the 60^th^ percentile of this distribution making them equal to the number of tri-loci fragments at the 60^th^ percentile.

The heatmaps in **Figures. 1K, 3B, 3C, 4B-4E** and **5** represent contiguous genomic regions. To observe interactions within these genomic locations we binned these regions accordingly so that each cell in the heatmap can be linked with two genomic locations on the x-axis (using two straight diagonal lines stemming from this cell). The intensity of the interaction between these genomic locations is represented by the color of the cell (**Figure 1K**). Specifically, the normalized intensity of interaction for each cell is given by log[1+#reads]/max(log[1+#reads]) where the denominator is associated with the cell with the highest intensity in the heatmap. Color-bars in **Figures. 1K, 3B, 3C, 4B-4E and 5** represent the range of the number of tri-loci events shown in each figure.

### RNAseq methodology

We used a cDNA sequencing protocol (adapted from http://wasp.einstein.yu.edu/index.php/Protocol:directional_WholeTranscript_seq) that preserves information about transcriptional direction. This allows more accurate determination of the structure and expression of genes. We also use a ribo-depletion step rather than the classic oligo-dT selection (targeting RNAs polyA) to purify out the ribosomal RNAs as many non-coding RNAs may not be polyadenylated. Paired-end reads were mapped to the mm10 genome using TopHat2 (parameters: --no-coverage-search --no-discordant --no-mixed --b2-very-sensitive -N 1).

## RESULTS

### Workflow for extracting multi-loci chromatin interactions using 4C-seq data

4C-seq generated with NlaIII as a primary restriction enzyme cutter is likely to capture short fragments due to the high frequency of cutting. Thus using this 4bp increases the probability that 3C products encompassing multiple interactions are identified. It is important to note that 4C reads which contain a secondary restriction enzyme and two or more additional loci are not necessarily indicative of multi-loci interactions since these constructs can arise from cross ligation of digested products derived from different cells. To improve detection of multi-loci events that originate from single cells, we only retain reads in which distinct loci are separated by sites associated with the first restriction enzyme. In other words, only fragments with two or more primary restriction enzyme sites are considered for identification of multi-loci interactions (**Figure 1G and H**). Using this strategy for alignment of multi-loci interactions, we were able to identify two or more loci that interact simultaneously with the bait region within a single cell.

The probability of obtaining multi-loci 4C fragments depends on read length. Indeed, the longer the read length the greater the probability of finding fragments with at least two primary restriction enzyme sites that are not interrupted by a secondary restriction enzyme site. Since reads representing four or more locus interactions are rare in experiments with 100bp reads, we will restrict our discussion to tri-loci interactions.

Our method complements existing 4C-seq pipelines, as it captures multi-loci interactions in addition to the standard pairwise interactions between the bait and other genomic locations. Furthermore, it allows us to differentiate between *bona fide* tri-loci interactions (bait-loci2-loci3) and mutually exclusive pairwise interactions (bait-loci2 or bait-loci3).

### Frequency and genomic distance associated with multi-loci interactions

To analyze multi-loci interactions we performed sixteen 4C-seq experiments using four different baits (3’Eκ, MiEκ, DSR and Eβ) located on chromosome 6 (**Figure 2A**) on *ex-vivo* derived cells from two lineages and four stages of mouse lymphocyte development: Double Negative (DN) and Double Positive (DP) T cells, Pre-(Pre-B) and Immature (Imm B) B cells. For the Eβ, 3’Eκ and MiEκ baits were designed within regions adjacent to each enhancer but not overlapping with sequences eliminated in knockout cells. All experiments were performed using NlaIII as the primary and DpnII the secondary restriction enzyme. We followed the workflow described above for analysis of each library and compared the frequency of multi-loci interactions across the sixteen experiments (**Figure 2B**). We observed that the frequency of tri-loci interactions is consistent between replicates for the same bait sequence and this is strongly dependent on the distribution of restriction sites around the bait. To understand this better, we counted the number of potential multi-loci fragments within 0.1Mb from the Eβ or 3’Eκ bait. Interestingly, in experiments using the Eβ versus the 3’Eκ bait we found more potential tri-loci interactions indicating that the genomic location of a bait may contribute to the probability with which multi-loci fragment interactions can be detected. In line with this finding we identified a significantly higher frequency of tri-loci interactions associated with the Eβ versus the 3’Eκ bait on the *cis* chromosome (P-value=0.005). (**Figure 2C**).

**Figure 2:**
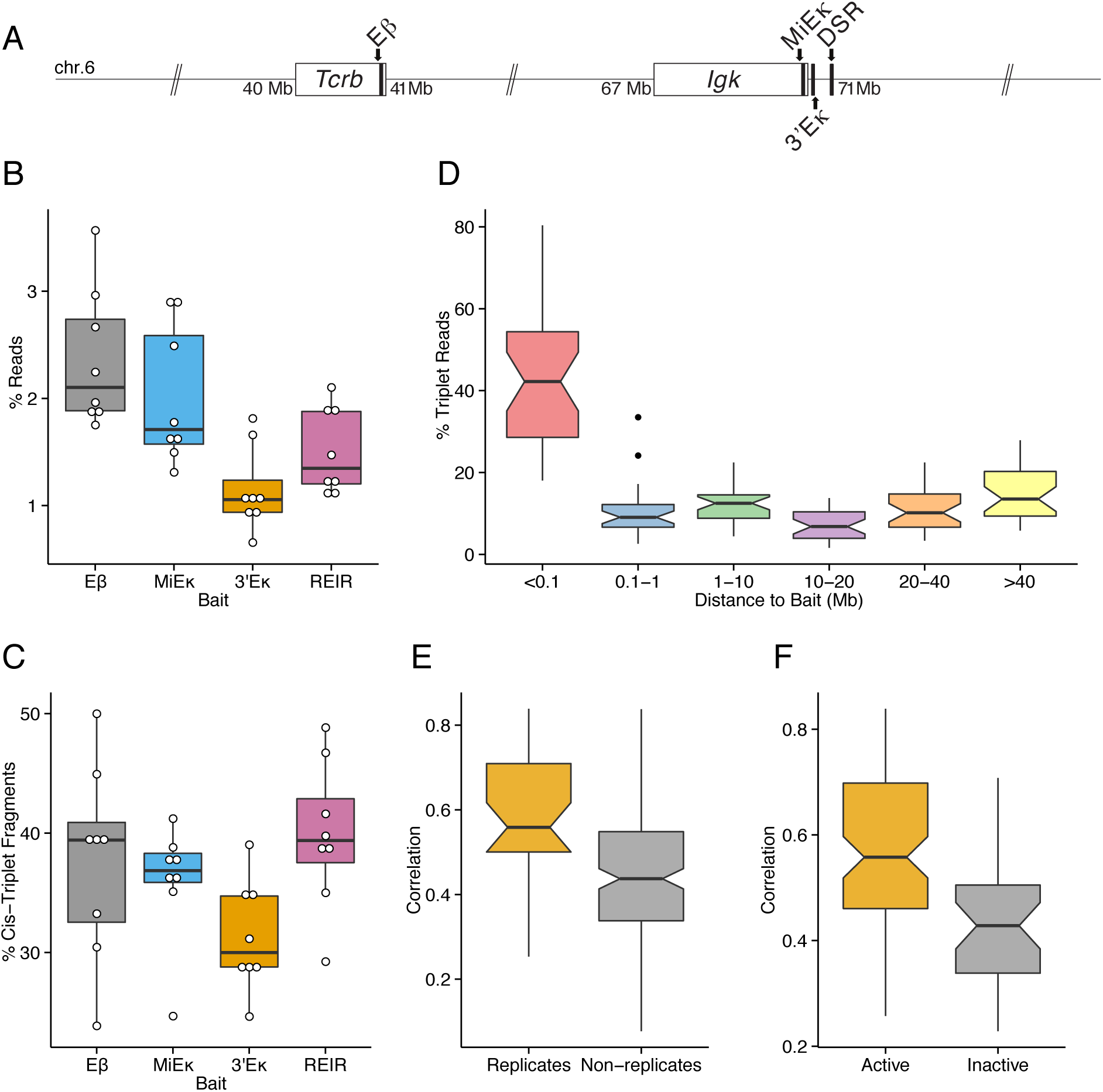
Frequency and genomic distance associated with multi-loci interactions. **A.** Scheme showing the location of the four 4C-seq baits on chromosome 6. **B.** Frequencies of tri-loci reads in 4C-seq experiments using baits Eβ, MiEκ, 3’Eκ, and DSR **C.** Frequencies of *cis* tri-loci fragments. **D.** Distribution of the frequencies of multi-loci fragments at different distances from the bait region. **E.** Correlation of pair-wise interactions between replicates and non-replicates. **F.** Correlation of interactions using active versus inactive baits.

We next, examined the genomic distance separating individual loci in tri-loci interactions (**Figure 2D**). In agreement with pairwise interactions detected in 4C-seq, we found enrichment of tri-loci interactions in regions located in close proximity to the bait. Specifically, 40% (median estimate) of tri-loci events occur within 0.1Mb from the bait. This indicates that regions in close linear proximity are more suitable for revealing simultaneous multi-loci interactions with the bait.

As a consistency test, we generated an interaction matrix, denoted as M1, whose columns represent twenty-four experiments involving two replicate experiments that use the baits MiEκ, 3’Eκ and DSR in four different cells types (DN, DP, Immature B, and Pre-B) and whose rows represent the intensity of tri-loci interactions within windows sizes of 500Kb across the genomic co-ordinates 68M-72M, the region surrounding these three baits on chromosome 6. For the Eβ bait, we generated a similar matrix, denoted as M2, whose columns represent eight experiments involving two replicate experiments in DN, DP, pre-B and Immature B cells and performed the same analysis to examine tri-loci interactions within window sizes of 50Kb across the genomic co-ordinates 40.8M-41.8M, the region surrounding Eβ on chromosome 6. We compared the tri-loci interaction matrix of any two experiments from the same bait and found that as expected correlations between replicates are significantly higher than correlations between experiments in two different conditions (**Figure 2E** P-value=6*10^−4^).

Since MiEκ, 3’Eκ, and DSR are active in B cells and Eβ and DSR are active in T cells we next compared the tri-loci interaction matrix of any two experiments where the bait is in an active versus inactive state. The correlations of tri-loci interactions of active baits are significantly higher than those of inactive baits P-value=6*10^−4^) (**Figure 2F**).

Taken together these findings are consistent with findings from other chromosome conformation capture experiments (11,18). The frequency of tri-loci interactions identified is dependent on the frequency of the restriction enzyme sites and the length of the reads. In addition, tri-loci interactions are detected more frequently in regions surrounding the bait and interactions are more reproducible if the bait is in an active versus inactive state.

### 4C-seq multi-loci interactions reflect known lineage and stage specific changes in *Tcrb* locus conformation

As a second consistency test, we separately checked the tri-loci interactions in *Tcrb* from a bait region with its associated enhancer, Eβ. For this we used window sizes of 50kb across the genomic co-ordinates 40.8M-41.8M surrounding the *Tcrb* locus on chromosome 6. We applied principal component analysis (PCA) to the interaction matrix M2 and found that the leading principal component (45% of the total variation) separates DN cells from DP and B cell subsets (**Figure 3A**). To understand these interactions in more detail we pooled replicates and compared the normalized tri-loci interaction intensity in DN, DP, pre-B and Immature B cells (**Figure 3B-E**). In DN cells where *Tcrb* undergoes recombination and the Eβ is in an active state we identified two local interacting clusters. Specifically, interactions are detected within and between the proximal diversity (D), joining (J) and constant (C), DJCβ1/2 region and distal variable Vβ domains, while the regions encompassing the inactive trypsinogen gene clusters have less contacts (**Figure 3B**). These data reflect known changes in locus conformation that occur in recombining DN cells: namely that Vβ genes interact more closely with each other and with the proximal DJCβ1/2 domain as a result of ‘locus contraction’ and that these interactions facilitate Vβ-DJβ rearrangement (21). However, from these analyses we now also identify simultaneous interactions within and between these two clusters. The absence of contacts within the trypsinogen clusters results either from looping out of this region or from deletion that will occur after Vβ -DJβ joining events. There are far fewer interactions detected in the Vβ cluster in DP cells, the developmental stage that follows on from rearrangement, (**Figure 3C**). This likely reflects the fact that many Vβ gene segments have been deleted as a result of rearrangement. In contrast, the absence of interactions across the *Tcrb* locus, particularly within the Vβ gene cluster in either pre-B or immature B is reflective of an absence of ‘locus contraction’ or inter-locus interactions in these cells (21) (**Figure 3D and E**). Taken together these data demonstrate the feasibility of our method in identifying known pairwise interactions involving the *Tcrb* enhancer and in revealing how these participate in tri-loci hubs.

**Figure 3:**
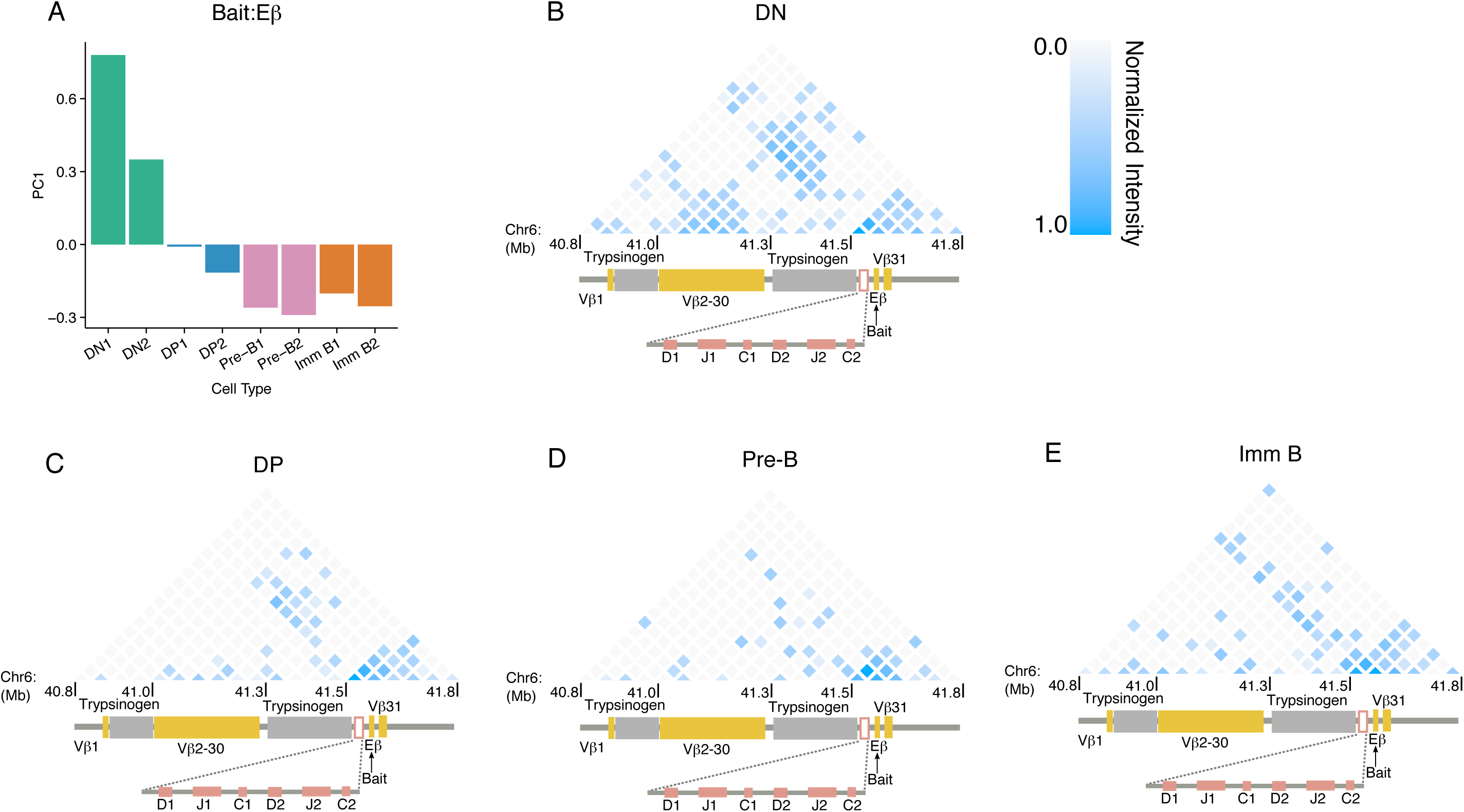
4C-seq multi-loci interactions reflect known lineage and stage specific changes in *Tcrb* locus conformation. **A.** The first principal component of pair-wise interactions in DN, DP, pre-B and Immature B cells using Eβ as bait. **B-E.** Heatmap for tri-loci interactions involving the Eβ bait in DN, DP, pre-B and Immature B cells across the *Tcrb* locus.

### 4C-seq multi-loci interactions reflect known lineage specific changes in *Igk* locus conformation

To investigate if interactions reflect lineage specific changes in locus conformation we checked tri-loci interactions across *Igk* from the MiEκ and 3’Eκ baits using window sizes of 200kb across the genomic co-ordinates 68M-72M on chromosome 6. For this we applied PCA to the interaction map to M1. The leading principal components (contributing 45.9% and 11.2% of total variation) indicate that the two B cell subsets (pre-B and Immature B) were significantly separated from the two T cell subsets (DN and DP). As expected we found no significant differences between experiments using the MiEκ and 3’Eκ baits (**Figure 4A**).

**Figure 4:**
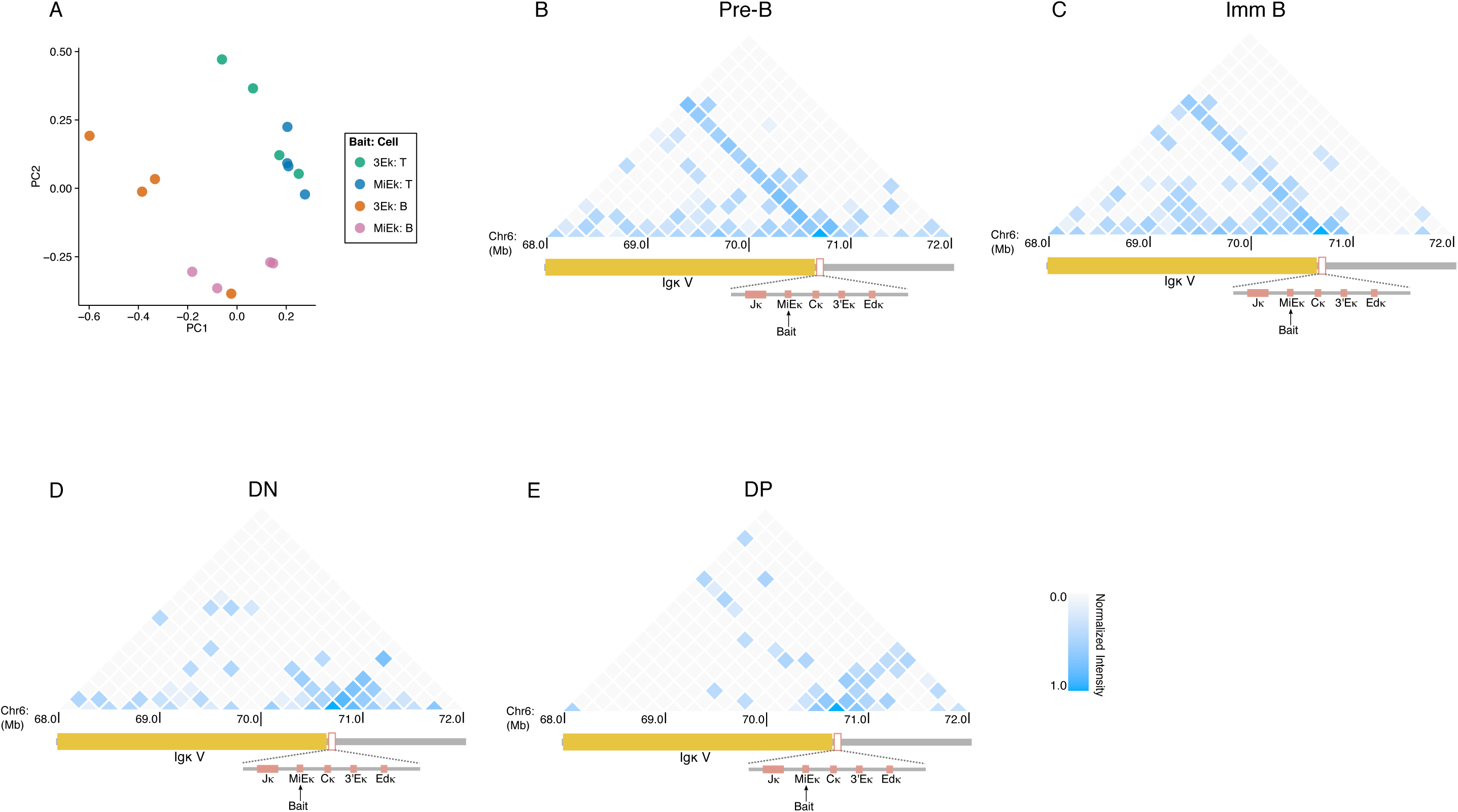
4C-seq multi-loci interactions reflect known lineage specific changes in *Igk* locus conformation. **A.** Scatterplot of the first and second principal components of pair-wise interactions of DN, DP, pre-B and Immature B cells using MiEκ and 3’Eκ as baits. **B-E**. Heatmaps for tri-loci interactions involving the MiEκ bait in pre-B, Immature B, DN and DP cells across the *Igk* locus.

Next, we pooled the replicates and displayed the normalized tri-loci interaction intensity in the four cell types, DN, DP, pre-B and Immature B for the MiEκ and 3’Eκ baits (**Figure 4B-E and Figure S2A-D, respectively**). Heatmaps for the MiEκ and 3’Eκ baits of the same cell type show similar interaction patterns in agreement with the PCA. There are several clustered regions in the heatmaps of the B cells consistent with *Igk* locus contraction occurring in recombining B cells (22).

Rearrangement of the *Igk* locus occurs predominantly via inversional as opposed to deletional rearrangement so the Vκ gene segments are largely retained within the locus following rearrangement and as a result we detect little change in the contacts across the region in immature B cells. This is in contrast to what we observed for the *Tcrb* locus in DP cells where rearrangement occurs via a deletional process. (**Figure 3C**). As expected we see fewer contacts along the *Igk* locus in T cells, however the enhancers interact more with regions outside of the locus at the 3’ end in these cells (**Figure 4D, E and Figure S1C, D**). Together these results demonstrate that our method can detect known lineage specific changes in locus conformation in addition to identifying simultaneous interactions that may influence the topology of antigen receptor loci and be important for V(D)J recombination.

### Interdependent tri-loci interactions at the 3’ end of *Igk*

The MiEκ and 3’Eκ enhancers are known to promote *Igk* recombination (23,24). These two enhancers interact with each other and another downstream *Igk* enhancer, Edκ that has no known function in recombination. To examine the relationship between the three enhancers we generated six heatmaps of tri-loci interactions using window size of 2.5kb. We pooled the reads from pre-B and Immature B cell 4C-seq experiments using MiEκ and 3’Eκ as baits in WT, MiEκ knockout, and 3’Eκ knockout conditions (**Figure. 5A-5F**). 4C-seq experiments with the 3’Eκ or MiEκ baits showed strong simultaneous local contacts between MiEκ, 3’Eκ and Edκ in WT B cells (**Figure 5A, 5D**). Interestingly, we found that the constant region Cκ located between 3’Eκ and MiEκ was looped out. Of note, we further show that deletion of either the MiEκ or the 3’Eκ not only interferes with tri-loci MiEκ, 3’Eκ and Edκ interactions in which they participate, but they also disrupt pairwise interactions between the two other enhancers in the hub (**Figure 5B, 5C, 5E, 5F, 5G, 5H**). Concomitant with these changes we also find a reduction in transcriptional output at these other partner loci (**Figure 5B**). These findings highlight the interdependent nature of multi-loci associations and their functional importance in gene regulation and provide new insight into chromatin organization as a whole.

**Figure 5:**
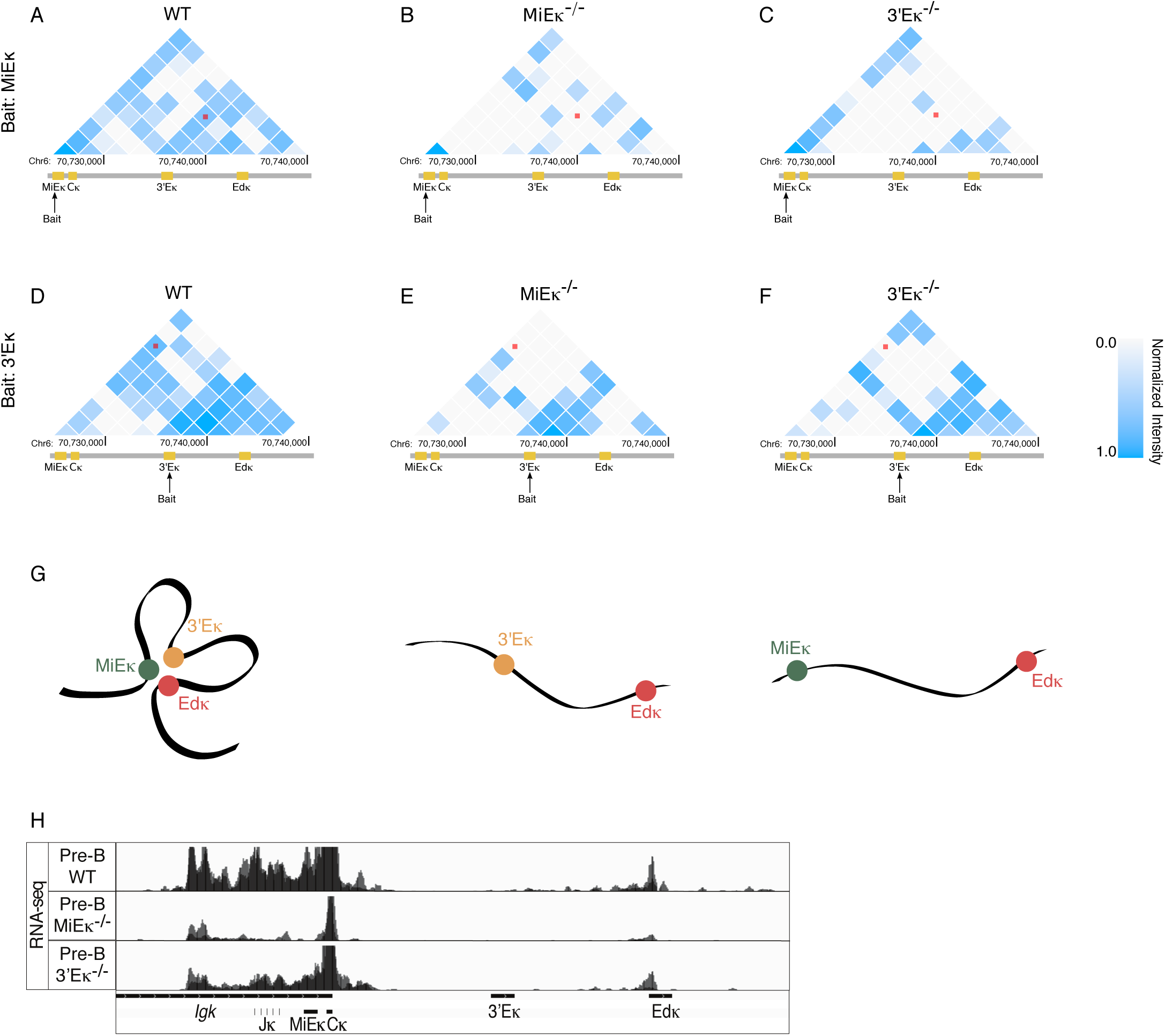
Interdependent tri-loci interactions at the 3’ end of *Igk.* **A-C.** Heatmap for tri-loci interactions involving the MiEκ bait in WT versus MiEκ^−/−^ and 3’Eκ^−/−^ B cells across the 3’ end of the *Igk* locus. Red dots mark interactions between the three enhancers. **D-E.** Heatmap for tri-loci interactions involving the 3’Eκ bait in WT versus MiEκ^−/−^ and 3’Eκ^−/−^ B cells across the 3’ end of the *Igk* locus.

## DISCUSSION

Although we know of many examples that underscore the importance of pairwise chromatin interactions in gene regulation (7-11), we have almost no information about the impact of multi-loci associations. Here we present a method that uses 4C-seq data to identify multi-locus interactions from the same cell. We demonstrate the feasibility of our method using 4C-seq data that identifies known pairwise interactions that occur within the *Tcrb* and *Igk* antigen receptor loci and additionally extract information that tells us which of these is involved in tri-loci associations. We further show that enhancer deletions not only interfere with tri-loci interactions in which they participate, but they also disrupt pairwise interactions between other participating partner loci. This suggests that multi-loci interactions have an unappreciated role in gene regulation and that our current models for how this works are oversimplified.

Recent Hi-C analysis and related approaches have uncovered the fact that the genome is organized into megabase size TAD structures (15-18). It is thought that one potential function of these highly conserved structures is to locally restrict the action of regulatory elements to loci contained within their boundaries. The identification of these new structures then opens up a number of important questions concerning gene regulation. For example, although we now know that promoters are not specifically tethered to specific enhancers in a one-to-one relationship (25), we do not yet know whether simultaneous interaction of enhancer elements with an individual promoter contributes to the regulation of individual genes (26,27). Our pipeline provides a tool to address this question.

Our methodology could also be used to examine the relationships between individual enhancer elements that make up ‘super-enhancers’ (28). As yet we do not know whether the individual enhancers that make up these elements are in contact and whether simultaneous interactions between component enhancers are important for their ‘super’ status. The analysis we have described here suggests that not only do component enhancers interact they can do so simultaneously. Moreover, these interactions can be interdependent such that if one contact is lost it impairs the contacts between other components in the hub. Finally, our pipeline provides a tool to determine if gene regulation involving multiple elements involves simultaneous interactions that are dictated by hierarchical relationships.

## ACKNOWLEDGMENTS

We would like to thank members of the Skok and Kluger labs for helpful suggestions and discussions. We would also like to thank NYU CHIBI and the NYUMC sorting and genome facilities for their contributions to this work. CP and VS, sorted developing lymphocytes from wild-type and enhancer knockout mice and generated the 4C-seq and RNA-seq data. TJ, YK, JS, RR, CP, VS, PR, RB and SB developed the idea for identification of tri-loci interactions and TJ implemented the pipeline. JS, TJ and YK wrote the manuscript with comments from all authors.

## FUNDING

PR is a National Cancer Center and American Society of Hematology Fellow. JAS is a Leukemia & Lymphoma Society (LLS) scholar. This work was supported by NIH grants R01 GM086852 (JAS and YK) and R01GM112192 (JAS).

**Figure S1:**
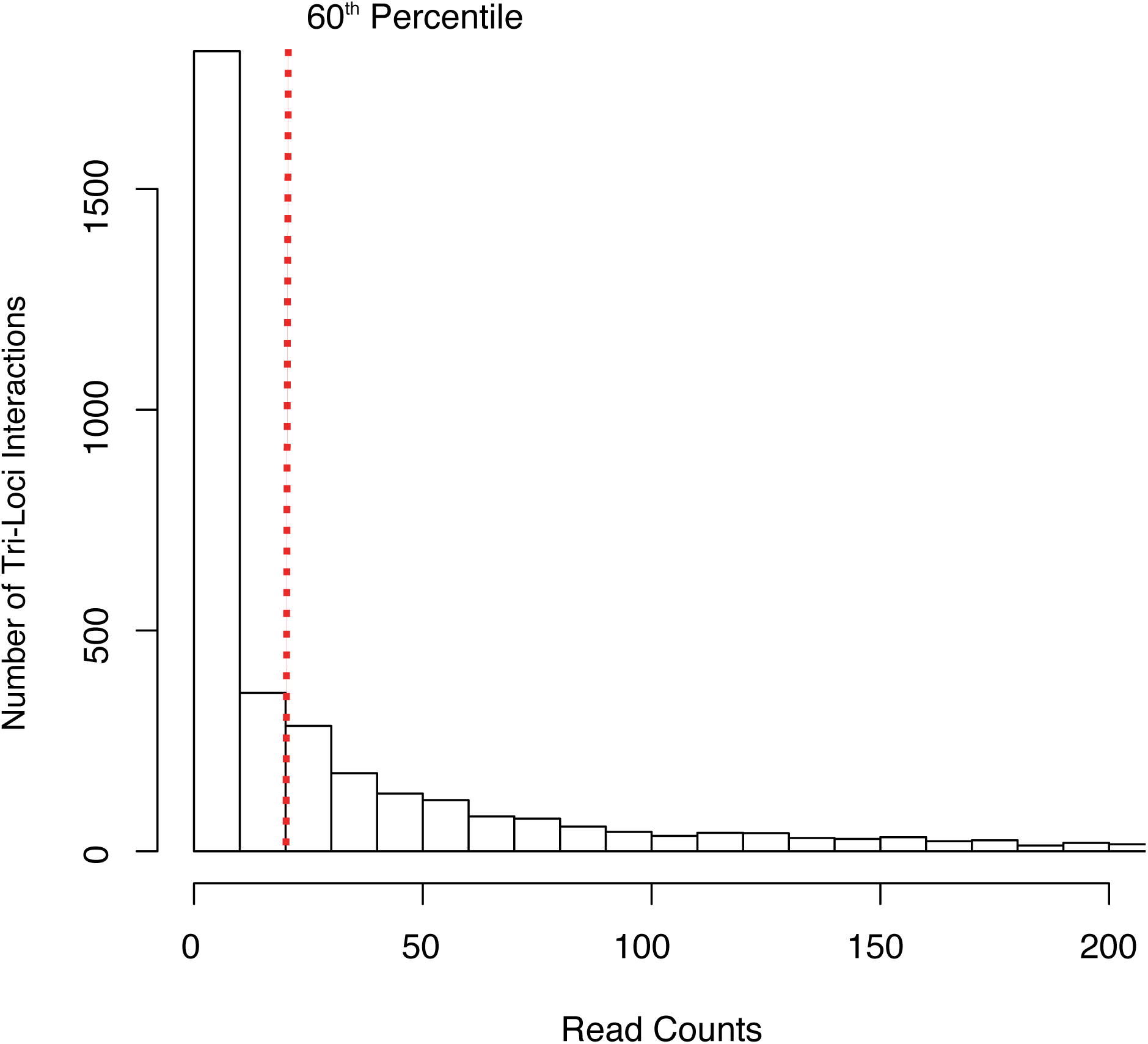
The frequency of distinct tri-loci interactions as a function of the number of identical tri-loci reads. To reduce the effect of PCR artifacts we capped the number of identical tri-loci fragments above the 60^th^ percentile making them equal to the number of identical tri-loci fragments at the 60^th^ percentile. For example, if the 60^th^ percentile line is positioned at the 4^th^ (x=4) bin of the histogram, each tri-loci read that is identically duplicated more than four times (and hence associated with bins with x>4) is counted as if it were sequenced only four times.

**Figure S2:**
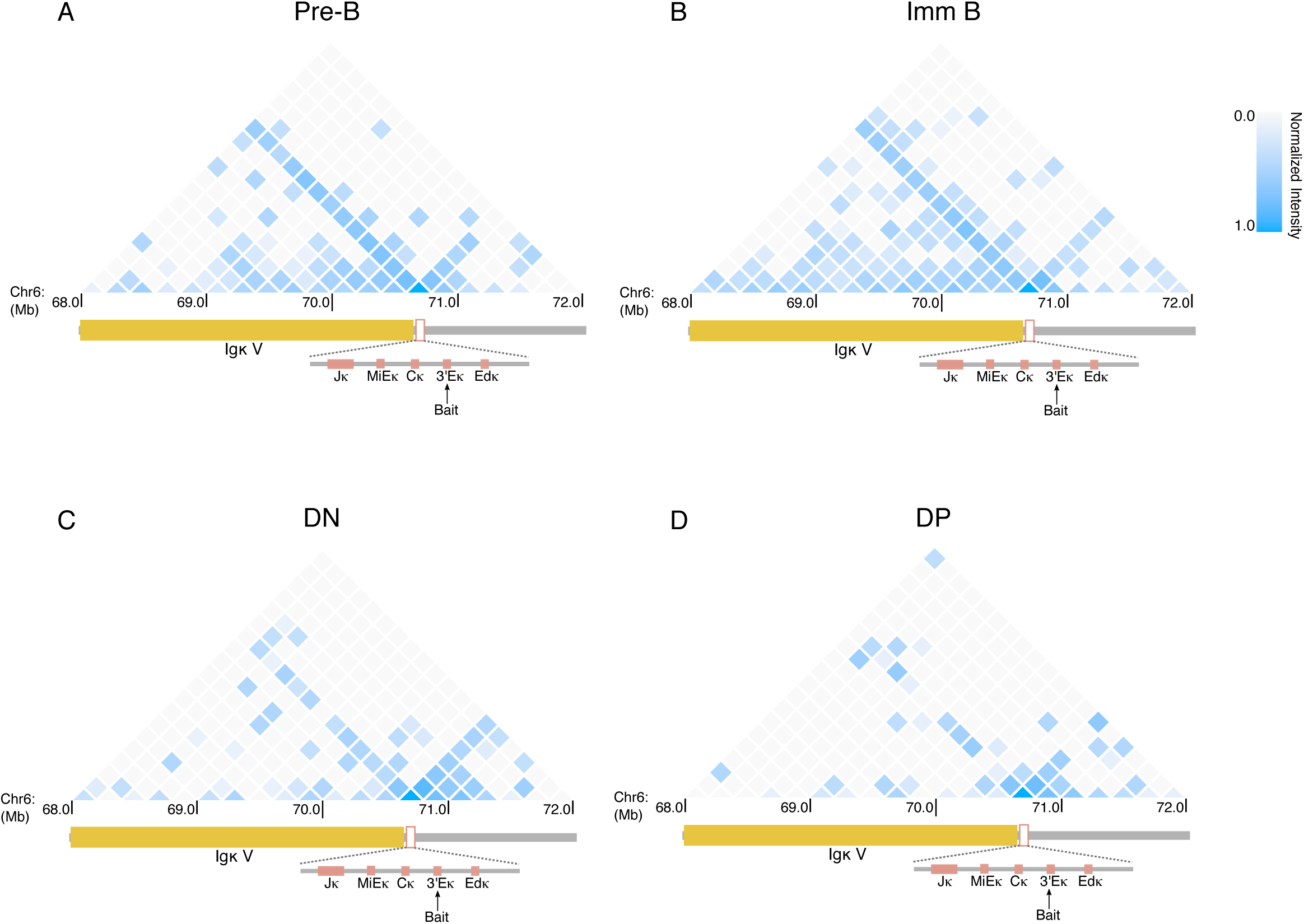
4C-seq multi-loci interactions reflect known lineage specific changes in *Igk* locus conformation. **A-D** Heatmaps for tri-loci interactions involving the 3’Eκ bait in pre-B, Immature B, DN and DP cells across the *Igk* locus.

**Supplementary Table:** Number of reads obtained for each 4C-seq experiment

| Experiments | Total Read Count |
| --- | --- |
| Bait:3Ek_DN_WT_rep1 | 16305102 |
| Bait:3Ek_DN_WT_rep2 | 38414105 |
| Bait:3Ek_DP_WT_rep1 | 14837187 |
| Bait:3Ek_DP_WT_rep2 | 32746291 |
| Bait:3Ek_ImmB_WT_rep1 | 13400974 |
| Bait:3Ek_ImmB_WT_rep2 | 49342558 |
| Bait:3Ek_preB_WT_rep1 | 12082764 |
| Bait:3Ek_preB_WT_rep2 | 37819428 |
| Bait:MiEk_DN_WT_rep1 | 14951357 |
| Bait:MiEk_DN_WT_rep2 | 8021581 |
| Bait:MiEk_DP_WT_rep1 | 11005056 |
| Bait:MiEk_DP_WT_rep2 | 8019384 |
| Bait:MiEk_ImmB_WT_rep1 | 11750140 |
| Bait:MiEk_ImmB_WT_rep2 | 8809675 |
| Bait:MiEk_preB_WT_rep1 | 24943586 |
| Bait:MiEk_preB_WT_rep2 | 10991332 |
| Bait:DSR_DN_WT_1 | 36997739 |
| Bait:DSR_DN_WT_2 | 11381608 |
| Bait:DSR_DP_WT_1 | 7419337 |
| Bait:DSR_DP_WT_2 | 10156868 |
| Bait:DSR_ImmB_WT_1 | 9062442 |
| Bait:DSR_ImmB_WT_2 | 10725219 |
| Bait:DSR_pre-B_WT_1 | 9171175 |
| Bait:DSR_pre-B_WT_2 | 13178411 |
| Bait:Eb_DN_WT_rep1 | 11884996 |
| Bait:Eb_DN_WT_rep2 | 31873206 |
| Bait:Eb_DP_WT_rep1 | 12687310 |
| Bait:Eb_DP_WT_rep2 | 51047901 |
| Bait:Eb_ImmB_WT_rep1 | 15371306 |
| Bait:Eb_ImmB_WT_rep2 | 45585596 |
| Bait:Eb_preB_WT_rep1 | 19530612 |
| Bait:Eb_preB_WT_rep2 | 55577812 |

